# The regulation of tDNA transcription during the directed differentiation of stem cells

**DOI:** 10.1101/108472

**Authors:** J. L Woolnough, D. A. Schneider, K.E. Giles

## Abstract

The human genome consists of 625 tDNA copies that encode 51 distinct isoacceptor families. Recent studies demonstrated that changes in chromatin structure during cellular differentiation can alter the expression of these tDNA. However, the mechanism by which tDNA can be differentially regulated remains unclear. Here we used the directed differentiation of pluripotent human embryonic stem cells (hESCs) towards the endoderm lineage as a model system to study the developmental regulation of individual tDNA. We demonstrated a significant change in the Pol III occupancy at 49 tDNA (22 reduced and 27 increased). The regulation of tDNA did not correlate with changes in TFIIIB or TFIIIC occupancy, H3K4me3, or H3K27me3 levels. However, tDNA that had an increase in Pol III binding were preferentially found within strong CTCF-COHESIN chromatin loops. The knockdown of either Ctcf or Rad21 in mouse tail tip fibroblasts had similar effects on changes in tRNA levels. We identified 7 isoacceptors that were differentially expressed during the directed differentiation of hESCs. The open reading frames of the ribosomal protein genes, which are translationally repressed during hESC differentiation, are enriched for codons that are decoded by these downregulated isoacceptors. Thus, translation efficiency during cellular differentiation may be affected by changes in tDNA regulation.

## Introduction

Transfer RNA are short, heavily modified, and highly structured RNA that serve as the essential adapter molecules during mRNA translation (1). These abundant RNA account for up to 15% of total cellular transcripts and are tightly regulated to compensate for changes in growth and stress conditions (2, 3). The transcription of tDNA requires the recognition of the gene-internal cis-regulatory boxes A and B being recognized by TFIIIC, which recruits TFIIIB to a site immediately upstream of the transcription start site (TSS). TFIIIB is then responsible for Pol III recruitment and the initiation of transcription (4). In higher eukaryotes, roughly half of the genomic tDNA are actively transcribed in any given cell and the genes which are active cannot be predicted by tRNA score (ie. strength of conserved sequence elements) (5). Genome-wide analyses have identified that many tDNA are transcribed in a cell-type specific manner (6, 7). The transcriptional activity of tDNA can correlate with local chromatin structure (e.g. H3K4me3, H3K27me3), Pol II localization, or the binding of transcription factors. However, there is currently no mechanistic understanding of how any of these factors contribute to cell-type specific changes in tDNA transcription.

tRNA abundance can regulate translation efficiency in prokaryotes and eukaryotes (8-10). Similarly, codon bias can be problematic when attempting to over-express mammalian genes in bacteria (11). However, the role of tRNA abundance in regulating translation efficiency as a normal mammalian gene expression program remains less clear. The population of tRNA is distinct between actively dividing and quiescent cells (12), and tRNA abundance correlates with codon usage among the *Drosophila melanogaster* proteome (13). Conversely, a computational analysis of the ability of tRNA from one cell-type to translate the mRNA from another concluded that there was no specific translational advantage conferred by cell-type specific populations of tRNA (14). Additionally, initial work investigating if dynamic tRNA expression was related to cell-type specific codon usage in eukaryotes, using ribosome profiling techniques, found no correlation (15-18). However, recent technological improvements in measuring tRNA abundance and translational efficiency have uncovered significant correlations between tRNA abundance and codon usage in the transcriptome of eukaryotes (19-21).

Here we assessed changes in Pol III occupancy during the directed differentiation of H9 hESCs into the endodermal lineage (22). This analysis identified the differential Pol III occupancy at 49 tDNA within 48 hours of ACTIVIN A treatment. These changes did not correlate with TFIIIB or TFIIIC occupancy, tDNA sequence, or local chromatin structure. ACTIVIN A did not regulate Pol III occupancy directly through the recruitment of SMAD2/3/4 to tDNA. However, tDNA that had an increase in Pol III occupancy were contained within strong CTCF-COHESIN chromatin loops. Further, the knockdown of Ctcf or Rad21 in mouse tail tip fibroblasts was sufficient to alter tRNA abundance. During the differentiation of hESCs we observed a significant change to 7 isoacceptors. The open reading frames of the ribosomal protein genes, which are translationally repressed during ESC differentiation, are enriched for codons that are decoded by these downregulated tRNA. This work suggests that chromatin structure can regulate individual tDNA and these changes impact translation during cellular differentiation.

## Results

### Pol III occupancy at tDNA is regulated during the differentiation of hESCs

To determine how Pol III occupancy was regulated during differentiation we first performed three replicates of ChIP-seq experiments for Pol III (Methods). To validate each experiment we used the MACs algorithm to identify peaks (23). We then identified the peaks that overlap with annotated tDNA (24). The overlap between each peak and a known tDNA is summarized via Venn diagram (Fig. 1A). The vast majority of all peaks overlapped a known tDNA in all three trials. We identified 291 tDNA that overlapped a Pol III peak in each of the three trials, which is consistent with published accounts that roughly 50% of all tDNA are transcriptionally active.

**Figure 1.**
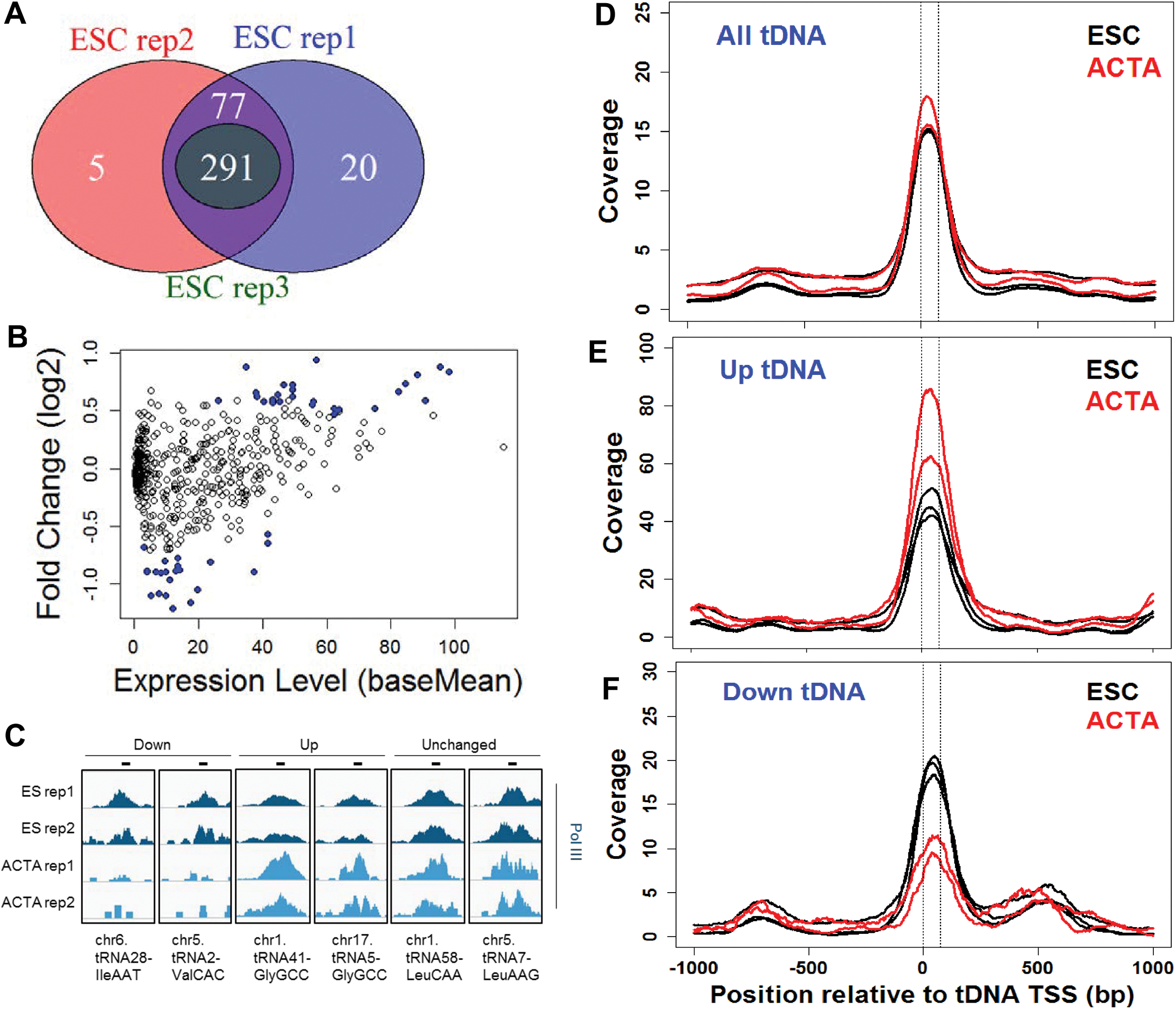
Pol III occupancy at tDNA changes during the differentiation of hESCs. (A) Three replicates of ChIP-seq for Pol III (RPC32: Santa Cruz #sc-21754) in H9 ESCs. Peaks of Pol III were called using MACS ver 1.4 (23), and the number of peaks that overlapped with annotated tDNA were determined using intersectBed (37). A Venn diagram shows the number of Pol III peaks at tDNA that overlap between the three replicates. (B) To understand how Pol III occupancy at tDNA could be regulated during differentiation we performed an additional two replicates of ChIP-seq after 48 hour treatment of H9 ESCs with ACTIVIN A (50ng/mL) to induce the hESCs to differentiate into endoderm. We then used the DESeq2 algorithm to quantify the occupancy of Pol III at each human tDNA in both hESCs and endoderm (39). An MA plot shows the fold change in Pol III occupancy at each tDNA (ACTIVIN A versus untreated H9 ESCs). The blue dots depict tDNA that underwent a significant change (p < 0.05, FDR ≤ 14%) in Pol III occupancy after 48 hours. (C) An IGV screenshot (41) of Pol III coverage at representative tDNA confirms the DESeq2 analysis. In each case the coverage was scaled according the sizeFactor determined by DESeq2. (D-F) A meta-analysis showing the coverage of Pol III (RPC32) within 1kb of each tDNA using the HOMER software suite (38). The tDNA were binned according to their change in Pol III occupancy: (D) unchanged, (E) upregulated, or (F) downregulated.

To determine if the occupancy of Pol III was regulated we repeated our ChIP-seq after treatment of H9 ESCs with ACTIVIN A for 48 hours, which induces differentiation into the endoderm lineage (25-27). A DESeq2 analysis was used to determine statistically significant changes to Pol III occupancy across all annotated tDNA (Methods). We detected 22 decreases and 27 increases in Pol III binding (Fig. 1B). These changes were confirmed through a visual inspection of the Pol III coverage across changed tDNA and six representative tDNA are shown (Fig. 1C). We further confirmed the DESeq2 results by performing a meta-analysis of the Pol III coverage within 1kb of the tDNA transcription start site (TSS) at either all tDNA, the upregulated tDNA, or the downregulated tDNA (Fig. 1D-F). In each case the meta-analysis was consistent with the results from DESeq2.

We next assessed if known tDNA cis-regulatory sequences could be responsible for the differential Pol III occupancy. This was accomplished by testing how many of each class of tDNA (unchanged, up, or down) contained an identifiable consensus sequence for either a TATA box, A- box, or B- box. A small percentage of all tDNA had a TATA box within 50 base-pairs upstream of the TSS (Fig. 2A, 19%). There was only a single “up” tDNA that had a TATA box compared to 27% of the “down” tDNA. Because the majority of tDNA did not have a TATA box we conclude that this cis-regulatory motif is unlikely to be responsible for the observed changes in Pol III occupancy.

**Figure 2.**
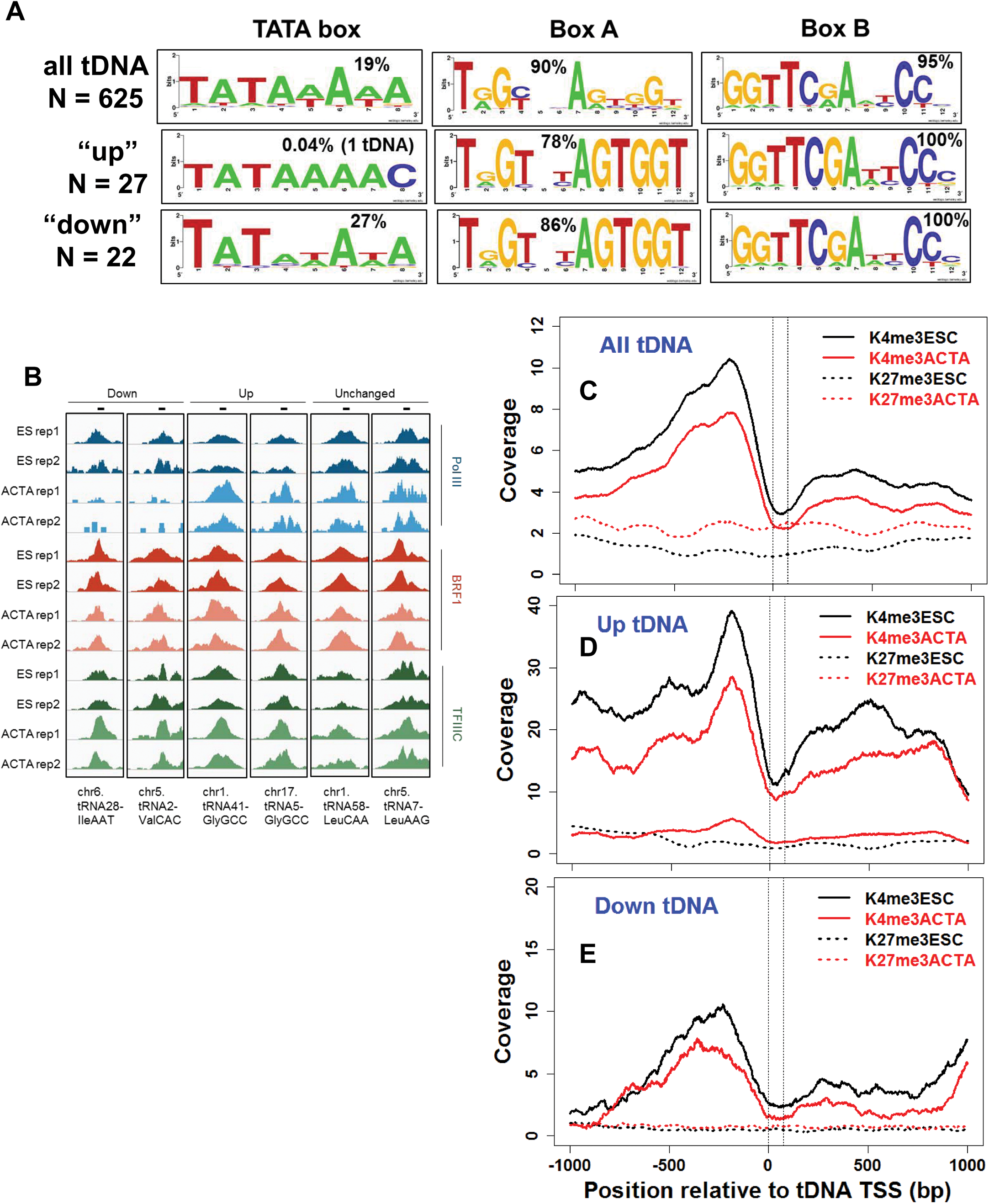
Changes in Pol III occupancy do not correlate with local cis-regulatory sequences, H3K4me3, or H3K27me3 levels. (A) HOMER was used to identify the consensus motifs for the TATA box, A box, and B box sequences within each tDNA bin (methods). The consensus sequences for the A and B boxes were adapted from previously reported genome-wide mapping of Pol III occupancy at tDNA (6). (B) We performed two replicates of TFIIIB (BRF1: Bethyl #A301-228A) and TFIIIC (TFIIIC-110: Santa Cruz sc-81406) ChIP-seq in untreated H9 ESCs and after 48 hours of ACTIVIN A treatment. Screenshots taken from the IGV of both replicates of TFIIIB and TFIIIC ChIP-seq. (C-E) The coverage of H3K4me3 and H3K27me3 within 1 kb of each tDNA was calculated using annotatePeaks within HOMER and plotted in Rstudio. The coverage for untreated and ACTIVIN A treated H9 ESCs was determined by re-analyzing the previously reported ChIP-seq datasets (GSM727586) (25). Normalization was carried out internally within annotatePeaks. The vertical dotted line indicates the position of the tDNA within each meta-analysis. The meta-analysis was performed on the set of all tDNA (C), the upregulated tDNA (D), and the downregulated tDNA (E).

We identified a B box within 100% of the differentially regulated tDNA, and there was no noticeable difference in the consensus sequence (Fig. 2A). Similarly, the identical A box consensus sequence was identified within 90% of all tDNA, 78% of “up”, and 86% of “down” tDNA. Thus, it is unlikely that these cis-regulatory motifs can explain the changes in Pol III occupancy.

We next examined the coverage of TFIIIB and TFIIIC at representative tDNA to determine if there was any correlation in occupancy at the individual tDNA level. This analysis clearly showed that where there were drastic changes in Pol III occupancy there was little or no difference in the occupancies of TFIIIB or TFIIIC (Fig 2B).

**Figure 3.**
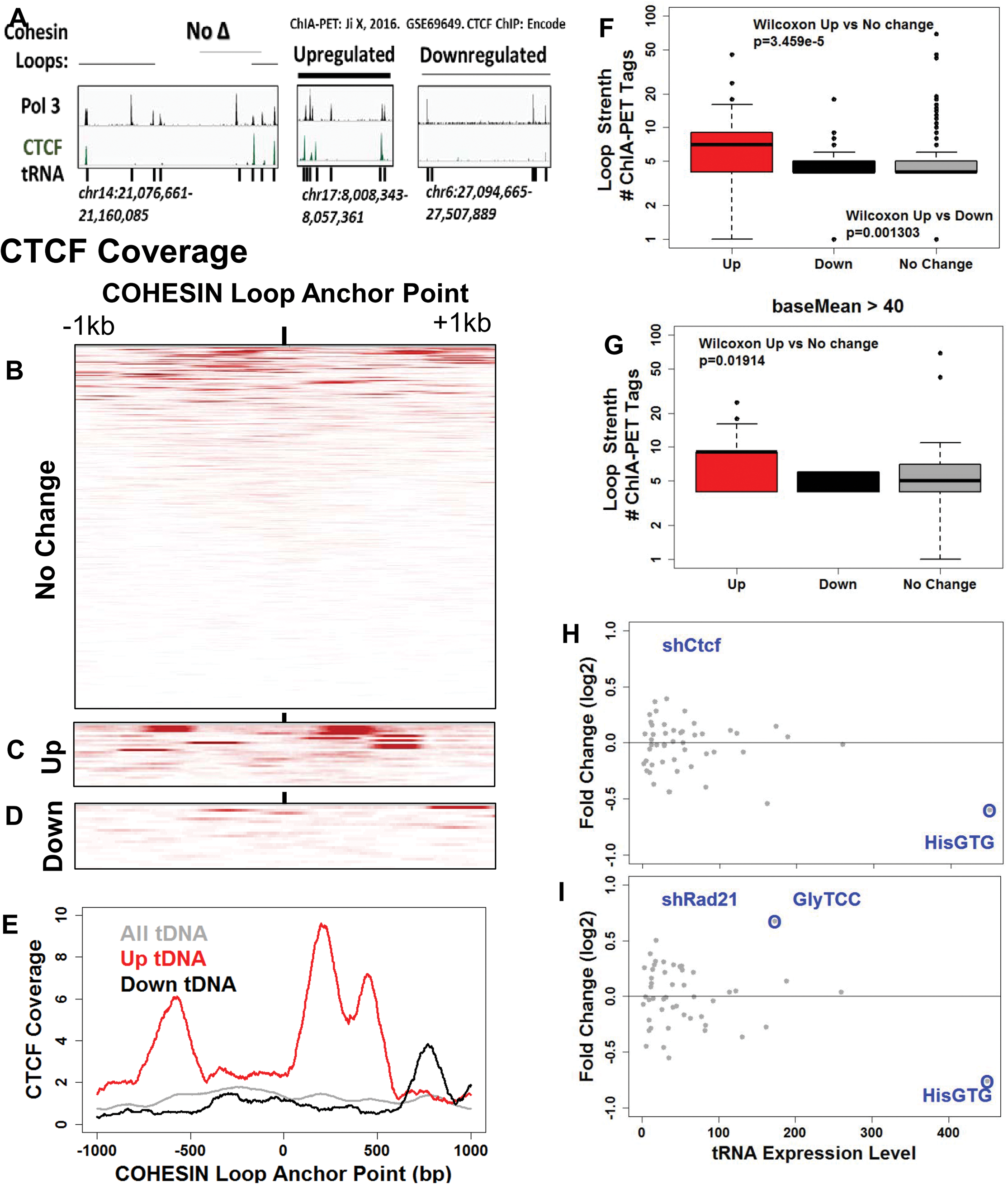
Chromatin structure regulates Pol III occupancy and tRNA levels. (A) A screenshot (IGV) showing the localization of Pol III (RPC32), CTCF (GSE69649), and tDNA for three genomic loci. The position of COHESIN loops, defined by ChIA-PET (30), are shown above each screenshot by a horizontal black bar. The thickness of each bar correlates with the number of ChIA-PET tags that span each loop. (B) The smallest loop that contains each tDNA was identified using BedTools intersectBed function. The coverage of CTCF within 1kb of the anchor points from each of these minimal loops was then determined using the –ghist command line argument with annotatePeaks. The numerical matrix was visualized using conditional formatting within Microsoft Excel, as previously described (34). The scale of the heatmap ranges from 0 counts (white) to the 99^th^ percentile of all counts (red). Three heatmaps were generated that depict CTCF coverage within 1kb of all tDNA (B), up tDNA (C), and down tDNA (D). (E) A meta-analysis demonstrates the average coverage across the genomic intervals shown in (B-D). (F) A box and whisker plot showing the distribution of loop strength for the three bins of tDNA (Up, Down, No Change). The loop strength was determined as previously reported as the number of ChIA-PET tags that span each loop (30). The upper and lower whiskers represent the 95^th^ and 5^th^ percentiles, respectively. The Wilcoxon rank sum test was performed to test if there was a statistically significant difference between the loop strengths of the “up” versus “no change” tDNA, and the “up” versus “down” tDNA. There was no significant difference between the “down” versus “no change” tDNA with respect to loop strength. (G) The distribution of loop strengths and statistical significance testing was done as described in (F), but with only tDNA that were occupied by Pol III with a baseMean greater than or equal to 40. (H,I) The levels of tRNA isoacceptors within control, shCtcf, and shRad21 mouse tail tip fibroblasts were quantified using a re-analysis of ribo-minus RNA-seq taken from GSE67516, as previously reported (31). The tags were aligned to each tDNA and then each tDNA was grouped into one of 51 isoacceptor groups and the tags were summed. Differential expression analysis was determined for shCtcf (H) and shRad21 (I) using DEseq2.

The presence of “activating” and “repressive” histone modifications have been shown to correlate with active and silent tDNA (28, 29). However, the functional relationship between local chromatin modifications and the transcriptional state of individual tDNA remains unexplored. Here, we used the differentiation of hESCs as a model to test the hypothesis that changes in H3K4me3 or H3K27me3 can facilitate changes in Pol III occupancy. This was carried out by a re-analysis of ChIP-seq data to quantify the levels of each modification within 1kb of each tDNA in control and ACTIVIN A treated hESCs (25). There was very little H3K27me3 across all tDNA, “up”, and “down” tDNA (Fig. 2C-E). The levels of H3K4me3 peaked upstream of the TSS and were decreased upon ACTIVIN A treatment, for all tDNA (Fig. 2C-E). The decrease in H3K4me3 signal may serve some regulatory purpose but is not likely to be responsible for the specific increases or decreases in Pol III occupancy because it was uniformly altered across all tDNA.

### Higher-order chromatin structure influences tDNA expression

The data presented in figure 2 indicated that local sequence and chromatin structure could not completely describe the differential occupancy of tDNA by Pol III in our system. This is consistent with the high level of conservation among various tDNA, making it difficult to discriminate between two tDNA based on local sequence. We therefore hypothesized that higher-order chromatin structure was an important factor in the regulation of tDNA. To determine the feasibility of this hypothesis we visualized CTCF binding in the vicinity of representative tDNA clusters (Fig. 3A). We observed that the binding of CTCF was greater in the vicinity of the “up” tDNA than for the “unchanged” or the “down” tDNA. To further investigate if this was a general trend we generated a heatmap of CTCF coverage within 1kb of the TSS for all tDNA, “up” tDNA, and “down” tDNA (Fig. 3B-D). The signal from each of these heatmaps was averaged and quantified in a meta-analysis (Fig. 3E). These analyses indicated that CTCF binding was much greater in the vicinity of “up” tDNA.

CTCF functions along with COHESIN to generate long-range chromatin loops that function to create independently regulated chromatin domains. To test if such loops could also be important in the regulation of Pol III occupancy at tDNA we determined the strength of previously reported chromatin loops that contained each tDNA (30). We visualized these loop strengths for the three representative clusters of tDNA via a horizontal black bar that connects two COHESIN loop anchor points (Fig. 3A). The thickness of the bar is scaled to be directly proportional to the number of reads that span the two anchor points. The loop that contained a representative “up” tDNA was significantly stronger than the loop that contained a representative “down” tDNA, as well as a cluster that contained only unchanged tDNA. To examine if this was a general trend we quantified the loop strength for each of the loops that contains an “up”, “down”, or “unchanged” tDNA (Fig. 3F). The median loop strength for those loops that contained an “up” tDNA was significantly greater than that for other tDNAs. To control for the fact that the “up” tDNA tended to have a higher overall Pol III occupancy than the “down” tDNA (Fig. 1B), we compared only tDNA that had a baseMean that was greater than 40. Despite this restriction, the median loop strength among the “up” tDNA was significantly higher than for other tDNA (Fig. 3G).

To determine if there was a functional relationship behind the correlation between loop strength and the regulation of Pol III occupancy we reanalyzed RNA-seq data from Ctcf and Rad21 (a Cohesin subunit) knockdown experiments in mouse tail tip fibroblasts (31). We utilized the ribo-minus RNA-seq data to quantify the levels of each tRNA isoacceptor (Methods), and performed DESeq2 to determine which of these 51 families were differentially expressed upon either Ctcf or Rad21 knockdown. Both knockdowns altered isoacceptor expression levels (Fig. 3 H,I). In particular, both Ctcf and Rad21 knockdowns caused a significant decrease in HisGTG. Although it remains to be determined how these knockdowns effect tDNA regulation, being that both caused a nearly identical change in HisGTG strongly suggests that the mechanism is related to the regulation of higher-order chromatin structure.

### Changes in tRNA abundance may contribute to the regulation of translation during cellular differentiation

To determine if changes in tRNA abundance can serve a regulatory function during cellular differentiation we quantified the changes in isoacceptor levels in control and ACTIVIN A treated hESCs. We identified 2 isoacceptors that were significantly decreased and 5 that were increased within 48 hours of ACTIVIN A treatment (Fig. 4A black dots). There were no significant changes after 1 hour (Fig. 4A, red dots).

**Figure 4.**
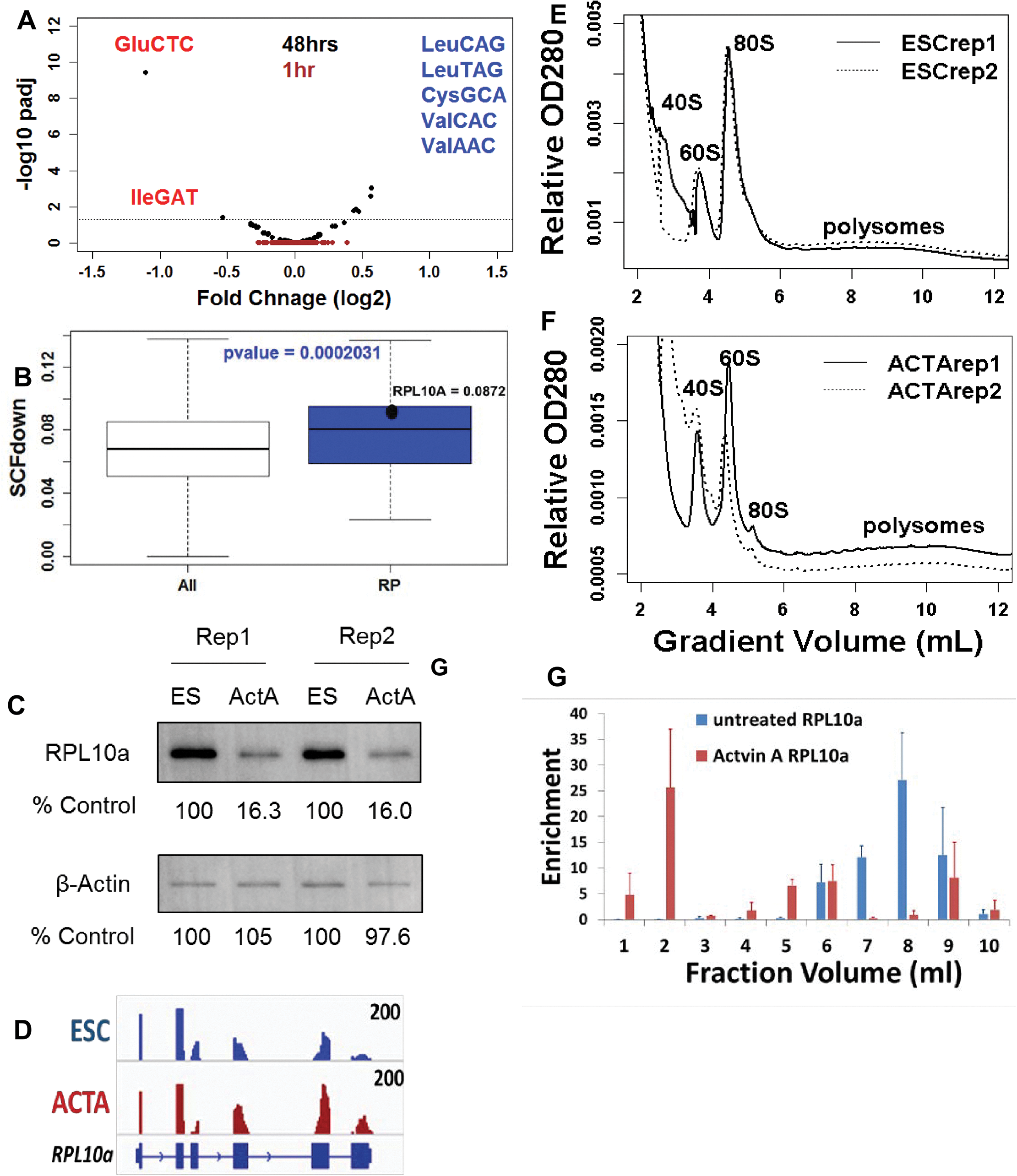
The regulation of tRNA isoacceptor abundance in human ESCs correlates with changes in translation efficiency. (A) A volcano plot showing the fold change in expression for each tRNA isoacceptor (n=51) after either 1 hour (red dots) or 48 hours (black dots) of ACTIVIN A treatment of H1 ESCs. The y-axis is the –log10 of the adjusted p-value, given by the Benjamini-Hochberg method. The isoacceptors that were significantly increased (padj < 0.05) are listed in blue, and those that were significantly repressed are listed in red. (B) The SCFdown for the codons (GAG, AUU, and AUC; see Methods) were calculated for a set of 52,748 transcripts and compared to that for the ribosomal protein genes (n = 115). The two distributions are visualized by a boxplot. The Wilcoxon rank sums test was used to determine a statistically significant increase in the SCFdown of the ribosomal protein genes (p = 0.0002). The SCFdown of RPL10A is indicated by a black dot. (C) A Western blot for both RPL10A and beta-actin from control and ACTIVIN A treated H9 ESCs. The protein levels for lane was calculated as a percentage of ES level and is shown beneath each blot. (D) IGV visualization of the RNA-seq counts at *RPL10a* in control and ACTIVIN A treated H9 ESCs, taken from (22). (E,F) Polysome profiles taken from control and ACTIVIN A treated H9 ESCs, as previously described (42). (G) Fractions were pooled from the polysome profiles and RT-QPCR was carried out to determine the migration of *RPL10A* mRNA in control and ACTIVIN A treated H9 ESCs. Each bar represents the average the two replicates for each cell type. The levels of the *RPL10a* mRNA were normalized to the migration of the 28S rRNA.

Previous studies have indicated that despite an overall increase in translation efficiency during ESC differentiation the ribosomal protein genes are translated less efficiently (15). This is consistent with the well-documented observation that rRNA synthesis, and therefore ribosome biogenesis, decreases during ESC differentiation (22, 32, 33). To investigate if there was potential connection between the down-regulation of specific tRNA isoacceptors and the repression of ribosomal protein gene translation, we created a metric that quantifies the fraction of each open reading frame that contains of a codon which is predicted to be decoded by one of the repressed isoacceptors. This metric, SCFdown, was determined by calculating the fraction of GAG, AUU, and AUC codons within each of 61,937 open reading frames. The median SCFdown value was significantly greater among the ribosomal protein genes than for the entire transcriptome (Fig. 4B).

We next wished to confirm the previous observations that specific ribosomal protein genes were translationally repressed in our model system. To that end, we demonstrated that RPL10A levels were reduced after 48 hours of ACTIVIN A treatment of H9 ESCs (Fig. 4C). There was no corresponding change in beta-actin. These changes in protein level were not driven by corresponding increase in mRNA levels (Fig. 4D) (22). We demonstrated that the changes in protein level were, at least in part, due to reduced translation efficiency by performing a polysome profile in both control and ACTIVIN A treated H9 ESCs (Fig. 4 E,F). We combined fractions of each polysome into 10 larger fractions and carried out RT-QPCR for *RPL10a* and *ACTB* mRNA. Fractions 1-5 roughly corresponded to the monosome fractions and fractions 6-10 to the polysomes. *RPL10a* was present exclusively within the polysomes within ESCs and increased throughout the monosomes upon ACTIVIN A treatment (Fig. 4G, top). Conversely, *ACTB* remained within the polysomes in addition to having a slight increase in the monosomes (Fig. 4G, bottom). These observations are consistent with translational repression for the ribosomal proteins during the differentiation of H9 ESCs with ACTIVIN A. We conclude that changes in tRNA abundance during differentiation may contribute to a novel form of translational regulation.

## Discussion

Large scale DNA sequencing projects, such as ENCODE, have revealed that many tDNA are located in regions of the genome that undergo cell-type specific changes in chromatin structure. These changes cause the differential expression of tDNA, which can alter the relative abundances of tRNA isoacceptor species. In many cases, tDNA silencing can be due to the formation of large regions of heterochromatin. However, here we show that significant changes to the occupancy of tDNA by Pol III can occur rapidly during cellular differentiation, in the absence of any drastic changes to chromatin structure. We have observed that Pol III binding to specific tDNA can be altered independent of TFIIIB or TFIIIC. Furthermore, these changes are not driven by known cis-regulatory sequences within the tDNA. We have also ruled out that these changes are driven by changes to H3K4me3 or H3K27me3 levels. However, the formation of higher-order chromatin structure, specifically chromatin loops anchored by CTCF and COHESIN have some role in regulating Pol III occupancy. We show that changes in tRNA isoacceptor levels occur during normal cellular differentiation and may contribute to the regulation of translation of specific classes of mRNA.

The regulation of Pol III activity has been well-studied. However, these studies have overwhelmingly focused on the bulk regulation of all tDNA during drastic cellular-wide changes, such as differentiation, stress, or cancer. There is very little understanding of how a specific tDNA could be precisely regulated. We hypothesized that this regulation would be related to the presence or absence of an upstream TATA box, or to different A- box or B- box sequences within the tDNA. However, we did not find any evidence that these regulatory regions are involved in the differential occupancy of Pol III. This does not mean that local cis-regulatory sequences are not involved. The ability of Pol II associated factors, including transcription factors, to bind upstream of many tDNA has been demonstrated. However, more work will be required to demonstrate the mechanistic roles for such transcription factors. In addition, although we have ruled out a role for H3K4me3 and H3K27me3, an exhaustive analysis of all known epigenetic marks must be carried out.

The intriguing observation that increased Pol III binding is correlated with strong chromatin loops suggests that genomic context is an important contributor to the regulation of tDNA. It is possible that these loops function to prevent the action of repressors of Pol III occupancy that are nearby, or that they facilitate the action of activators of Pol III occupancy. Future studies that delete specific CTCF binding sites in conjunction with swapping the genomic loci of “up” and “down” tDNA will be required to further understand the role of chromatin structure.

The observation that certain groups of transcripts, such as those for the ribosomal protein genes, are enriched for codons that are decoded by decreased tRNA isoacceptors is consistent with a recent report that demonstrated an inverse correlation between tRNA abundance and the ribosome dwell time at specific codons (21). Abundant isoacceptors entire the A site of the ribosome more rapidly and lead to more rapid translation. Conversely, a rare isoacceptor would cause the ribosome to stall and possibly result in lowered protein levels. Our data suggests that changes in tRNA abundance may facilitate such a mechanism during cellular differentiation.

## Methods

### Cell Culture and Drug treatment

H9 ESCs were obtained from WiCell as per agreement 15-W0341. ES cells were cultured and passaged as previously described (22). For ACTIVIN A cells, ES cell medium was replaced with RPMI with B-27 and ACTIVIN A (Sigma # SRP3003) at a final concentration of 50 ng/ml for 48 hours.

### ChIP

ChIP was performed as previously described (34, 35) but using the following antibodies: Pol III (Santa Cruz #sc-21754), BRF1 (Bethyl #A301-228A), and TFIIIC (Santa Cruz sc-81406). Samples were sequenced on an Illumina Hi-seq by either HudsonAlpha Genomics Services Laboratory or the UAB Heflin Center.

### Bioinformatics

All ChIP-seq data was processed through the following pipeline. FASTQ reads were first filtered using the fastx tool “fastq_quality_filter –q 20 –p 80”, which selects only reads that have at least 80% of its bases with a qual score greater than or equal to 20. These filtered reads were then collapsed to eliminate potential PCR duplication bias using fastx_collapser. The resultant fasta file was aligned to hg19 using bowtie2. The SAM file was converted into a BAM file and then BED file using Samtools “view” and bedTools “bamToBed”, respectively (36, 37). The bed files were used, in conjunction with our previously published input file from H9 ESCs (GSE76586), to call peaks using the MACS algorithm, using an FDR of 5% (23). The peaks were then intersected with the list of 625 tDNA from hg19 (taken from tRNAscan (24)). The bed files were also used as the inputs for the annotatePeaks script within HOMER for both meta-analysis and heatmap generation (38). Differential occupancy was determined using DESeq2 (39). The count tables were determined using intesectBed to identify all the ChIP-seq tags that originated within one of the 625 annotated tDNA. We randomized the sample names from the three trials of untreated H9 ESC and two trials of ACTIVIN A to directly estimate the false discovery method, as previously described (40). Briefly, every iteration that compared three of the five samples to the other two were created and DESeq2 was performed on each. We set a significance threshold of alpha = 0.05 and compared the number of false positives (each result with p < 0.05 in random trials) with the total number of significance calls (each result with a p < 0.05 in the actual comparison of H9 ESC versus ACTIVIN A). This method demonstrated a positive false discovery rate of 14% at alpha = 0.05. Transfer RNA isoacceptor abundance was determined by first aligning and counting the reads at each tDNA using bowtie. The tDNA were then grouped according to their isoacceptor (n=51) and the counts from each gene within that family were summed. This resulted in a count matrix with 51 rows, which was then input into DESeq2 to determine differential expression.

### Data acquisition

The Pol III, TFIIIC, and TFIIIB ChIP data from figures 1 and 2 are available in GSE94418. The histone ChIP data from figure 2 was reanalyzed from GSM727586 (25). The Cohesin ChIA-PET data from figure 3 was reanalyzed from GSE69649 (30). CTCF ChIP-seq is available from the ENCODE project as file wgEncodeBroadHistoneH1hescCtcfStdAlnRep2.bam. The isoacceptor analysis from figure 3 was reanalyzed from GSE67516 (31). The isoacceptor analysis in figure 4 was reanalyzed from GSE41009 (27). All processed files and accession numbers are deposited in the Gene Expression Omnibus: GSE94418.

